# Altered Attention Network Dynamics in Resting-State fMRI of Individuals with High Smartphone Addiction Scores

**DOI:** 10.64898/2026.07.26.740792

**Authors:** Harrison Watters, Aida Abdul Rashid, Subapriya Suppiah, Theodore LaGrow, Jaroslav Hlinka, Shella Keilholz

**Affiliations:** Institute of Computer Science of the Czech Academy of Sciences, Prague, Czech Republic; Weber State University, Ogden, UT, United States; Center for Diagnostic Nuclear Imaging, Universiti Putra Malaysia, Serdang, Malaysia; Wallace H. Coulter Department of Biomedical Engineering, Georgia Institute of Technology and Emory University, Atlanta, GA, United States

**Keywords:** Smartphone addiction, attention networks, resting state fMRI, dynamics, default mode network, attractor states, quasi-periodic patterns

## Abstract

Altered attention network activity has been consistently reported in recent neuroimaging literature of smartphone addiction and problematic smartphone use. However, many functional imaging studies have relied on traditional static functional connectivity analyses, which cannot capture time-varying or recurrent dynamics of attention networks. Given the importance of temporal and spatial dynamics for attentional processing, we applied two complementary dynamic functional connectivity approaches: functional connectivity-based attractor networks (fcANN), which characterize transitions between discrete attractor states related to internal-external context and action-perception dynamics, and cPCA-based quasi-periodic pattern (QPP) analysis, which captures phase relationships between cortical networks including the default mode network, dorsal attention network, and frontoparietal control network.

In a resting-state fMRI dataset comparing participants with High and Low SAS-M scores (SAS-M: Smartphone addiction scale-Malay version questionnaire), we observed reduced DAN participation within subject-specific cPCA-derived QPP-like components, along with context-dependent differences in DMN-attention-network relationships across attractor states. The most consistent effects emerged during transitions between internal and external attractor states, where subjects with high smartphone addiction (SPA) scores showed greater DMN dominance relative to attention-related networks. Overall, these findings provide preliminary evidence that smartphone addiction may be associated with altered large-scale attentional dynamics at rest

## 1. Introduction

People commonly use smartphones for 3-10 hours per day in high income economies (McCoy, 2020; Brodersen et al., 2022; Marciano & Camerini, 2022; Coyne et al., 2023; Nurbaiti, 2023; Tomczyk & Selmanagic Lizde, 2023). Assuming 8 hours of sleep per night (optimistic), this means many humans now spend more than half of their waking life engaged with smartphones; and about half of that on social media (Coyne et al., 2023; Tomczyk & Selmanagic Lizde, 2023). The shift towards obligate digital attention accompanies significant post-industrial changes in work-related physical activity, with many office and screen-oriented workers reaching sedentary screen time upwards of 15 hours per day (Levine, 2015; Jain et al., 2023).

These abrupt digital changes to human lifestyle are associated with numerous adverse health effects on sleep, metabolism, vision, and overall physical health (Nakshine et al., 2022), as well as elevated risk of psychiatric disorders such as depression, suicidal ideation, anxiety, narcissism, and attention deficit hyperactivity disorder (Vahedi & Saiphoo, 2018; Giordano et al., 2019; Kim et al., 2019; Ksinan et al., 2021; Brodersen et al., 2022; Nakshine et al., 2022). The behavioral correlates of digital overload, specifically smartphone addiction, are generally bleak. These behavioral effects raise an important question regarding their neural correlates. In terms of historical context, human brains evolved over several million years of foraging and hunting in small, tightly knit, communities - markedly different from the modern information landscape (Porzio, 2020; Burr, 2025). Naturally, there is significant interest in the *cognitive* cost of the devices, such as smartphones, which are shaping our modern social environment (Korte, 2020).

Neurofunctional changes associated with internet and smartphone addiction (SPA) are now widely reported (Lin et al., 2022; Anbumalar & Binu Sahayam, 2024). Predictably, association networks involved in cognition, executive function, and goal-directed external attention are among the most consistently implicated (Wang et al., 2019; Lee et al., 2021; Han & Kim, 2022; Kwon et al., 2022; Schmitgen et al., 2022; Áfra et al., 2024). However, cortical networks are shaped by mechanisms of plasticity throughout the lifespan, tuning connectivity to reflect environmental demands (Soares et al., 2013; Posner et al., 2014; Swingler et al., 2015; Power & Schlaggar, 2017; Klein et al., 2024). Lifelong neural plasticity complicates interpretation of whether smartphone-related neurofunctional changes should be viewed as maladaptive, adaptive, or context-dependent. While there is evidence that smartphone use can impair cognition and working memory (Ward et al., 2017; Stone, 2020; Akkurt & Aksoy, 2025), others argue that media-multitasking can improve attention (Kobayashi et al., 2020). Perhaps our brains are simply doing what they do best: adapting. In experienced videogamers for example, task-related activation in attention networks is altered, but the changes are associated with *improved* reaction times in visuospatial tasks (Jordan & Dhamala, 2023), and even short-duration gaming is linked to improved visual attention and executive function in elderly subjects (McCord et al., 2020). However, such improvements in performance may plateau, as video game experience improves multitasking only up to intermediate levels of difficulty and does not appear to protect against the effects of heavy media multitasking (Cardoso-Leite et al., 2016). In other words, plasticity can only compensate for so much multitasking.

Unlike video game experience, which often involves goal-directed problem solving, it is unclear whether extensive smartphone use, particularly with regards to social media, has any adaptive benefit (León Méndez et al., 2024). There is emerging evidence that SPA leads to maladaptive changes mimicking general features of addiction, such as volume changes in anterior cingulate cortex (ACC), a key region in motivation and will-power, as well as cue-related dopaminergic activity (i.e. craving) in the nucleus accumbens (NAcc), a primary reward center of the brain (Parvizi et al., 2013; Schmitgen et al., 2020; Abdul Rashid et al., 2021; Henemann et al., 2023), and changes to precuneus activity, related to impaired attention (Nasser et al., 2020). Excessive screen time has also been associated with reductions in cortical grey matter volume, with some authors suggesting possible implications for neurodegenerative risk later in life (Manwell et al., 2022).

While task- and resting-state functional MRI (rs-fMRI) studies have identified SPA-related alterations in attention and executive-control networks, most of the aforementioned work has relied on traditional static functional connectivity (FC) measures, in which relationships between regions of interest (ROIs) are averaged across an entire scan. Such approaches provide useful estimates of overall network organization, but do not directly characterize time-varying fluctuations or temporal phase relationships between networks (Keilholz et al., 2017; Bolt et al., 2022; Meyer-Baese and Watters, 2022). Because attentional processes fluctuate over time and across cognitive contexts, dynamic FC approaches may provide complementary information regarding large-scale network coordination in SPA. Indeed, there is evidence that phase information (temporal offset between network activations) might be more related to cognitive state than average activation or functional connectivity (Song et al., 2026).

To capture brain dynamics in a dataset of resting-state SPA subjects, we combined two approaches: functional connectivity-based attractor networks (fcANN) and quasi-periodic pattern (QPP) analysis. The attractor network approach, implemented by Englert and colleagues, captures dynamic brain states as free-energy minimizing attractor basins analogous to Hopfield Networks (Hopfield, 1982; Englert et al., 2025). The fcANN approach was found to be robust across datasets, sites, and acquisition parameters to reliably characterize four dynamic brain states in resting-state scans corresponding to two interpretable axes: internal vs external context, and action vs perception states (Englert et al., 2025). This approach offers the advantage of capturing task-like brain states in resting-state time series. Using their approach on a resting-state dataset of autism spectrum disorder (ASD), for example, they were able to detect alterations to sensory and default mode network (DMN) areas consistent with perceptual changes in ASD. Applying QPP analysis offers complementary insight, whereby cycles of anticorrelated activity between task-negative/task-positive regions of the brain, can be detected across imaging modalities (Majeed, 2011; Thompson et al., 2014; Xu et al., 2025). QPPs occur on various timescales, but are typically described as infraslow (∼0.05-0.1 Hz) cycles, meaning swells of network activity, lasting ∼20 seconds in group-averaged time series (Yousefi et al., 2018; Maltbie et al., 2022). While QPPs are robust across task and rest states, QPP network fluctuations appear flexible based on task demand (Watters et al., 2025). QPPs are largely driven by activity in the DMN, which is implicated in mind wandering, cognition, and memory recall (Raichle, 2015). DMN activity is typically antiphase with the dorsal attention network (DAN), which flexibly interacts with other networks such as the ventral attention network (VAN) and visual networks (VIS) to mediate externally oriented or goal-directed attention (Corbetta & Shulman, 2002; Vossel et al., 2014; Rajan et al., 2021; Zhao et al., 2022). The DAN in particular is a natural target of interest in SPA given its role in distractor suppression (Lanssens et al., 2020). Lastly, the frontoparietal network (FPN), sometimes called the executive control or control network (Witt et al., 2021) is associated with executive function and information selection (Scolari et al., 2015; Marek & Dosenbach, 2018), and has demonstrated reduced activity during distractor tasks among smartphone addicted subjects, with corresponding failures of executive control (Han & Kim, 2022). The balance of activity and phase offset between cortical networks such as the DMN, DAN, and FPN during QPP events appears to be important to attention and task performance (Seeburger et al., 2024; Seeburger et al., 2026). There is additional evidence that QPP dynamics are disrupted in disorders of sleep and attention (Abbas et al., 2019; Daley et al., 2025). Applying QPP analysis in SPA is thus a natural extension of the method.

Each of these dynamic approaches provides complementary information about large-scale brain organization. The attractor-network framework characterizes recurring brain states and transitions between attentional contexts, including internal versus external orientation and action versus perception-related states. QPP analysis, in contrast, characterizes recurring spatiotemporal patterns and temporal phase relationships between cortical networks, including interactions involving the default mode and attention systems. Applied together, these approaches permit examination of how large-scale network phase relationships vary across brain states and transitions during resting-state activity.

This study presents an exploratory application of combined attractor-state and QPP analyses to characterize resting-state network dynamics in relation to smartphone addiction. Using fcANN, we examined whether SPA was associated with altered occupancy and transition structure across attractor-defined attentional states, including internal versus external and action versus perception-related modes of network organization. Using QPP analysis, we further examined whether SPA was associated with altered phase relationships among large-scale resting-state networks, particularly involving the DMN, DAN, FPN, and related sensory-attentional systems. Finally, by relating QPP dynamics to attractor-defined states and transitions, we examined whether large-scale network phase relationships varied systematically across different modes of resting-state brain organization.

## 2. Materials and Methods

### 2.1 Participants and Dataset

Resting-state fMRI data included 45 participants organized by the original study team into High SAS-M (n = 21) and Low SAS-M (n = 24) groups. These supplied SAS-M group assignments were retained for the present study. Five participants were excluded during preprocessing quality control because of excessive head motion or gross spatial-registration failure (three High SAS-M and two Low SAS-M), yielding a final sample of 40 participants (High SAS-M: n = 18; Low SAS-M: n = 22). The resulting score distributions were consistent with the previously validated SAS-M screening threshold of >98 for identifying individuals at elevated risk of problematic smartphone use (Ching et al., 2015; Figure 1A). Note that the SAS-M is a screening instrument rather than a clinical diagnostic assessment, thus, the groups are referred to as High and Low SAS-M in the manuscript. All participants were right-handed, had normal or corrected-to-normal vision, and reported no history of neurological or psychiatric disorders. The groups did not differ significantly in sex distribution (two-sided Fisher’s exact test, p= .339); sex and age counts are shown in Figure 1.

**Figure 1:**
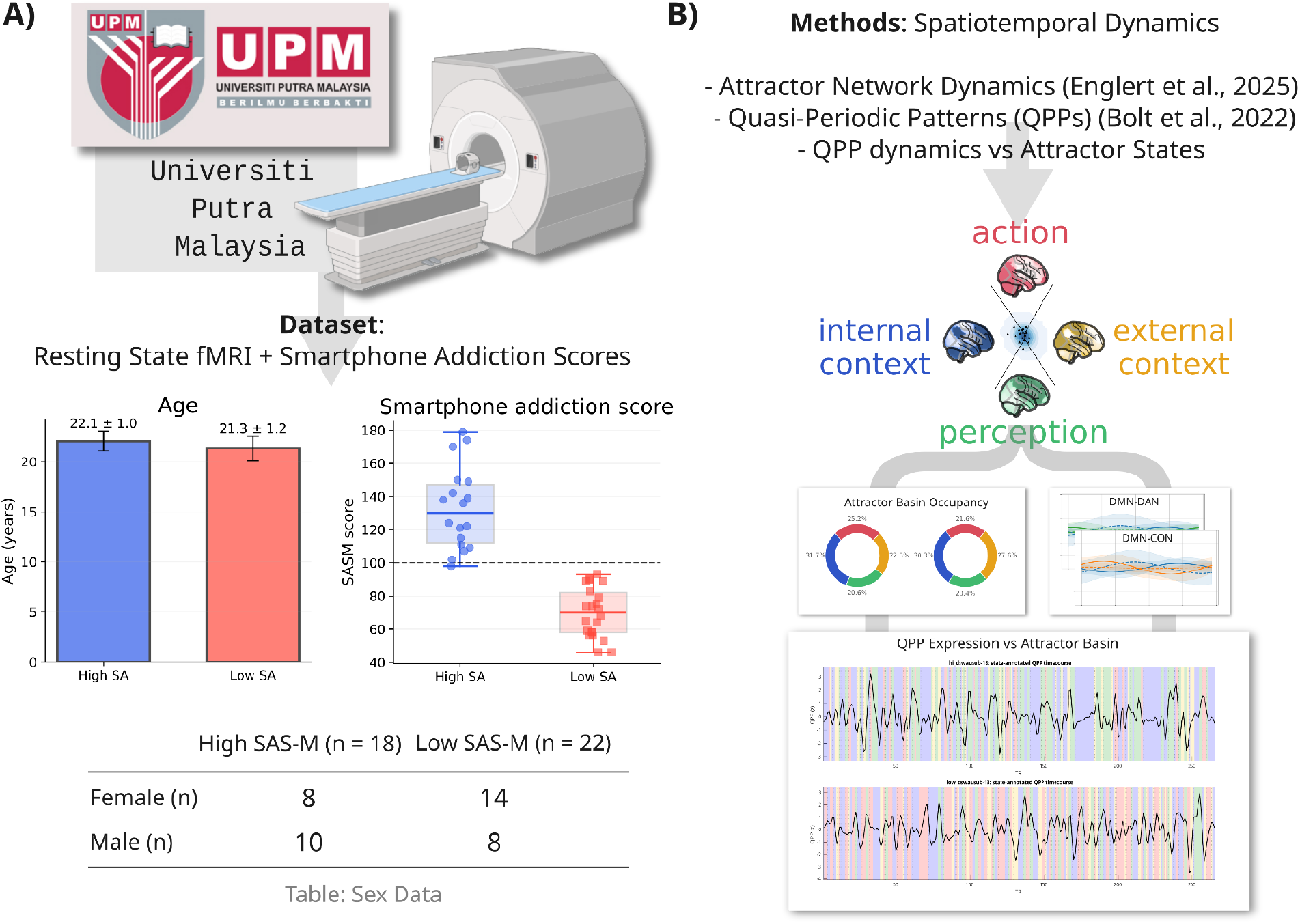
Study Overview. **A)** Dataset summary showing participant demographics and smartphone addiction scores (SAS-M), with subjects stratified into High and Low smartphone addiction (SA) groups. Bar plots indicate mean age and SAS-M scores with standard deviation, and table summarizes sex distribution across groups. **B)** Schematic overview of the analysis pipeline. Resting-state fMRI data were processed using two complementary dynamic analysis approaches: functional connectivity-based attractor networks (fcANN) and quasi-periodic pattern (QPP) analysis. Attractor analysis identifies discrete brain states along internal-external and action-perception axes, while QPP analysis captures recurring spatiotemporal patterns of large-scale network activity. Combined analyses examine QPP expression and network dynamics within and across attractor states. (MRI cartoon in panel A created with BioRender.com).

The original study was approved by the Ethics Committee of Research Involving Human Subjects of Universiti Putra Malaysia. All participants provided written informed consent. The present study constitutes a secondary analysis of previously acquired data.

### 2.2 MRI Data Acquisition

MRI data were acquired on a 3T scanner (PRISMA, Siemens, Erlangen, Germany).

Structural MRI scans were acquired with the following parameters: High-resolution T1-weighted magnetization-prepared rapid gradient echo (MPRAGE) sequence. Repetition time (TR) of 2300 ms, echo time (TE) of 2.27 ms, and inversion time (TI) of 1100 ms. Images were collected in an ascending sagittal orientation with 160 slices, a field of view (FOV) of 256 × 256 mm², matrix size of 256 × 256, isotropic voxel resolution of 1 mm.

Resting-state functional images were obtained using a gradient-echo echo-planar imaging (EPI) sequence sensitive to blood-oxygen level dependent (BOLD) contrast. The phase encoding direction was anterior-to-posterior. Functional acquisition parameters included a repetition time (TR) of 3000 ms and an echo time (TE) of 30 ms, with 256 TRs, giving a total scan length of ∼ 13 minutes per subject. Whole-brain coverage was achieved with 34 slices of 3 mm thickness, a field of view (FOV) of 220 × 220 mm², and a voxel size of 2.3 × 2.3 × 3 mm. Whole-brain functional images were collected during resting-state with participants instructed to remain still with their eyes closed but not fall asleep, and not engage in any specific task.

### 2.3 Resting-State Preprocessing

Functional MRI data were preprocessed using the CONN functional connectivity toolbox (Whitfield-Gabrieli, S., and Nieto-Castanon, 2012) implemented in MATLAB, which relies on SPM-based routines. Preprocessing followed a standard pipeline broadly comparable to that used in the original study, with minor differences in implementation.

Preprocessing steps included: Slice-timing correction to account for temporal offsets between slices, realignment (motion correction) using rigid-body transformation, coregistration of functional and anatomical images, segmentation of anatomical images into gray matter, white matter, and cerebrospinal fluid (CSF), normalization to the MNI-152 standard space, and spatial smoothing. Denoising (in CONN) included: regression of motion parameters, regression of physiological noise using principal components from white matter and CSF signals (CompCor approach), and linear detrending and temporal filtering (band-pass, 0.008-0.09 Hz)

These preprocessing steps are broadly equivalent to those used in the original study (which employed SPM-based preprocessing), ensuring comparability of the resulting time series.

All time series were parcellated to BASC-122 atlas space before analysis (Bellec, 2010). This 122 functional parcel atlas was selected to match the process used by Englert and colleagues for their Attractor Network approach, which was optimized on the same atlas (Englert, et al., 2025).

### 2.4 Comparisons and Correction

Group comparisons were conducted using two-sided Wilcoxon rank-sum tests (equivalent to Mann-Whitney U tests) and subject-level permutation tests, as specified for each analysis. Multiple comparisons were controlled using the Benjamini–Hochberg false-discovery-rate procedure where stated, with q < 0.05 considered significant. Effect sizes for between-group comparisons were quantified using Cohen’s d and, where reported, Cliff’s delta. Rank-sum and permutation-derived p-values are identified separately. Uncorrected results are presented only for exploratory interpretation and should be considered hypothesis-generating.

### 2.5 Attractor State Detection

Attractor states were identified using the *connattractor* framework (Englert, et al., 2025, PNI Lab), which implements a Hopfield neural network-based model of whole-brain functional connectivity dynamics. This approach uses a pretrained embedding derived from a large multi-site dataset, in which canonical brain states are defined as stable attractors in an energy landscape learned from functional connectivity patterns. This embedding was derived from an independent dataset (by Englert and colleagues) and was not trained on the present sample. Full methodological details and validation of this framework are described in the original study (Englert, et al., 2025), which demonstrated that these attractor states generalize across datasets, acquisition protocols, and temporal resolutions.

For the present study, preprocessed resting-state fMRI time series (BASC-122 parcellation) were projected into this pretrained attractor embedding using the default template provided by the connattractor package. At each time point (TR), the whole-brain activity pattern was evaluated within the Hopfield network to compute its corresponding energy and to determine its proximity to canonical attractor states. Each time sample was then assigned to the nearest attractor using connattractor functions *get_attractors_per_timesample* and *label_att_states,* resulting in a discrete time series of attractor labels for each participant. The original template defines multiple attractor states; for interpretability and consistency with prior work by Englert and colleagues, these were remapped to four canonical states reflecting two major cognitive axes (internal context, action, external context, and perception). For a sense of the cognitive axes, see Figure 2A.

**Figure 2:**
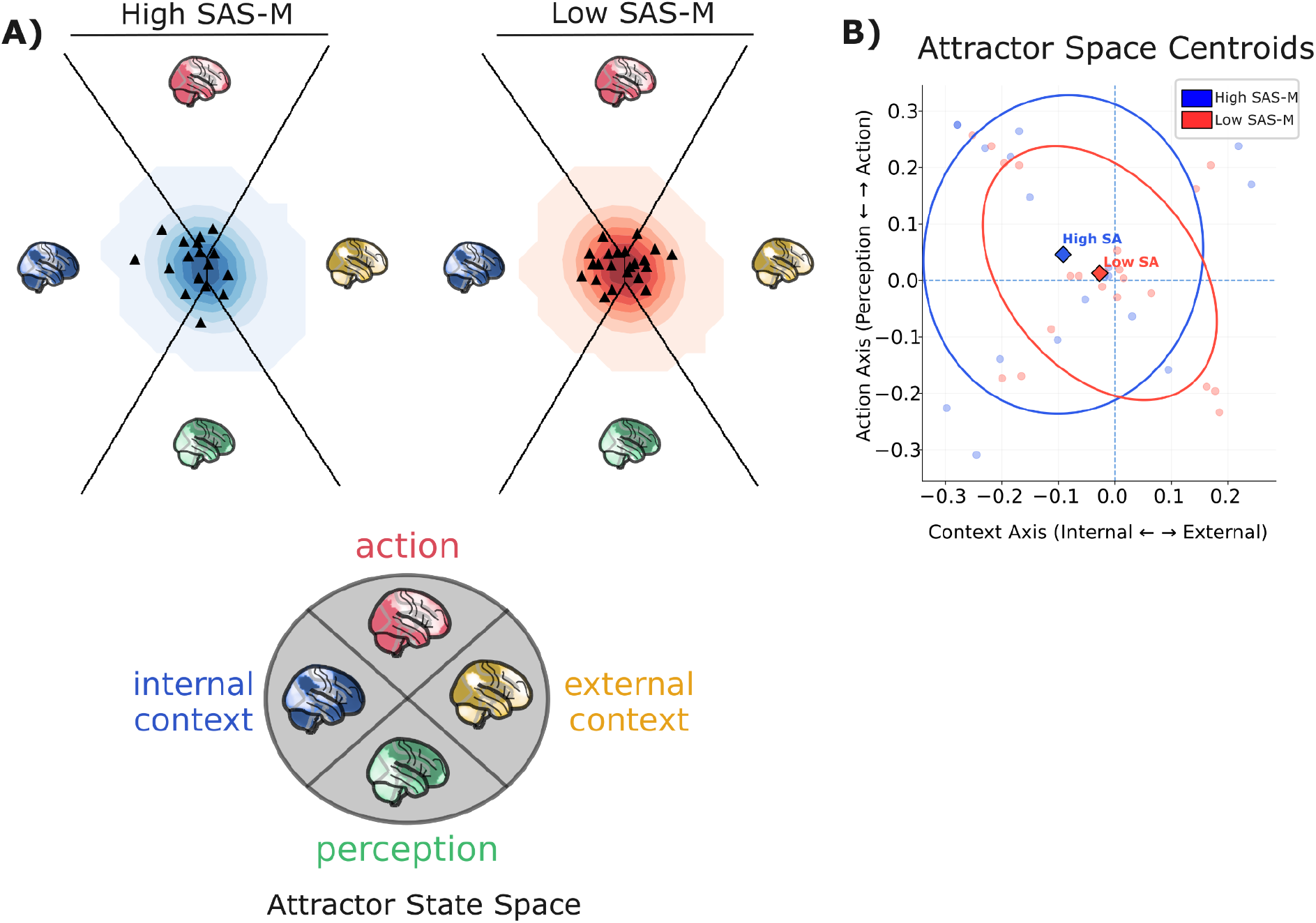
Attractor state-space density plots between groups. Whole-brain activity time series were projected into a shared two-dimensional space defined by the fcANN embedding. The horizontal axis represents the internal-external context gradient, and the vertical axis represents the perception-action gradient. **A)** State-space density maps show the distribution of time points for the High and Low SAS-M groups. Darker regions indicate greater time-point density. Black triangles indicate individual participants’ mean positions. **B)** Subject-level mean positions are displayed at an expanded scale to show variation near the origin. Diamonds indicate group centroids, ellipses represent approximately 1.5-standard-deviation covariance contours, and dashed lines mark the origin. The High SAS-M group showed greater descriptive dispersion and a small shift toward the internal-context and action directions; these visual differences were descriptive and were not subjected to formal inferential testing.

### 2.6 Attractor State Metrics

Following attractor state assignment, summary metrics were derived at both the time point and subject levels (Figure 3). For each participant, the fraction of time spent in each attractor state (state occupancy) was computed, along with entropy and switch rate. These state labels and associated metrics were subsequently used to examine relationships between large-scale brain network dynamics, quasi-periodic patterns (QPPs), and behavioral measures.

**Figure 3:**
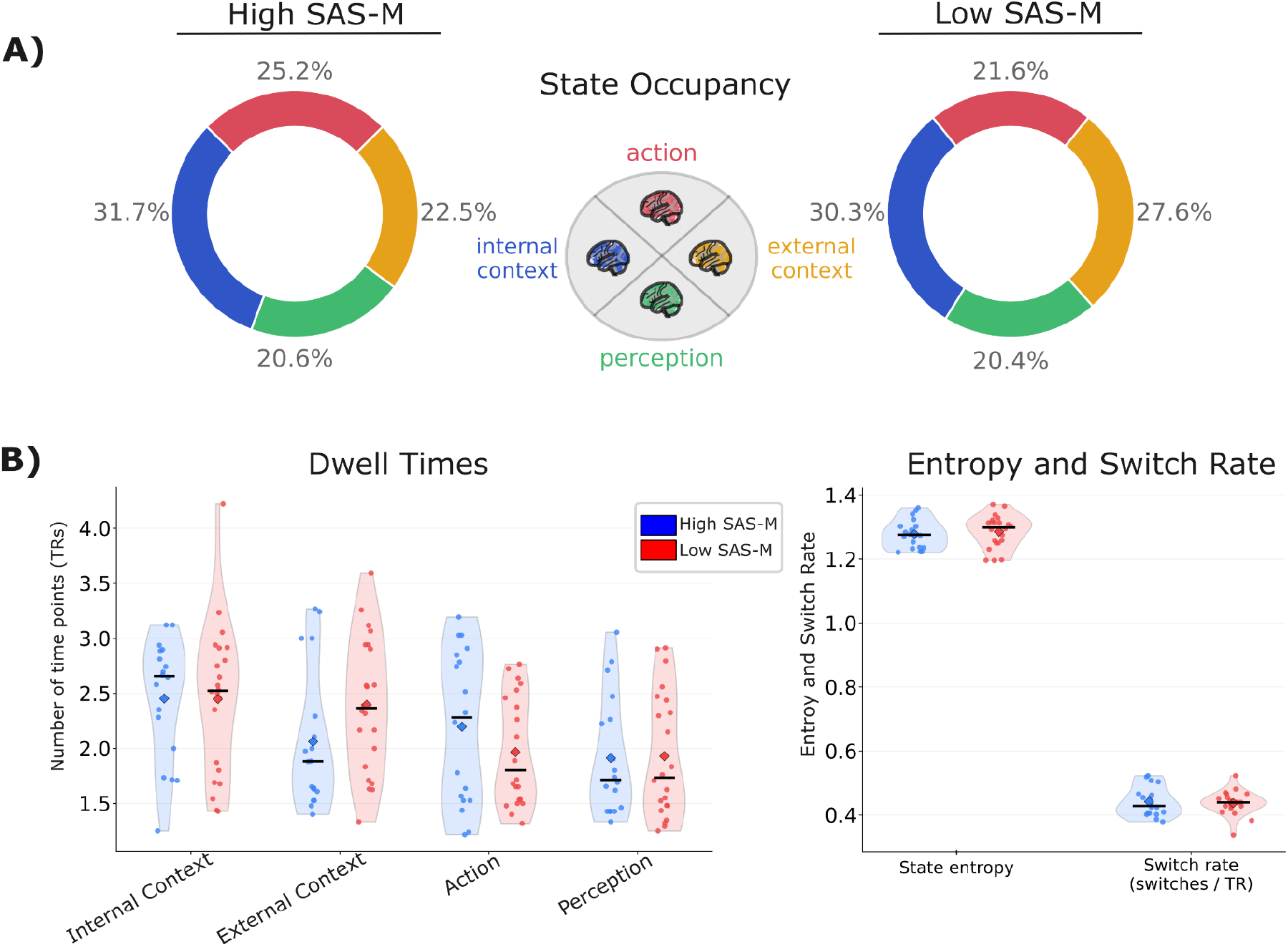
Attractor State Dynamics. **A)** Fraction of time (TRs/time points) spent in each attractor basin; internal - external, action - perception by group, High SAS-M = High smartphone addiction scores, Low SAS-M = low smartphone addiction scores. **B)** Additional attractor dynamic metrics including: mean dwell time within (from left to right) internal context, external context, action, and perception states, and in bottom right the entropy and switch rate (changes of state per TR). The four-state occupancy profiles (A) did not differ significantly between groups in the group label omnibus permutation test (p= .454; 10,000 permutations). Subject-level group comparisons of dwell time, entropy, switch rate, and axis metrics were assessed using two-sided Mann-Whitney U tests.

For attractor-state summary metrics (Figure 3), group differences were assessed at the subject level using two-sided Mann-Whitney U tests comparing High SAS-M (n = 18) and Low SAS-M (n = 22) participants for state fractions, context/action-axis summary scores, mean dwell times, state entropy, and switch rate. For descriptive interpretation, effect sizes were quantified as Cohen’s d and Cliff’s delta. Levene’s test (center = median) was additionally computed to assess heterogeneity of variance between groups for each metric. These analyses were performed on one value per subject for each metric.

To assess overall state occupancy in the donut plots (Figure 3), we performed an omnibus permutation test on the subject-wise occupancy vectors, using 10,000 random label permutations to obtain the null distribution.

### 2.7 Attractor State Specific Regional Activity Comparisons

State-specific regional activity was quantified by averaging parcel-wise BOLD time series within each subject and attractor state (Figure 4), yielding one mean value per parcel (BASC-122), per state, per subject. Group differences (High SAS-M vs Low SAS-M) were evaluated independently within each attractor state.

**Figure 4:**
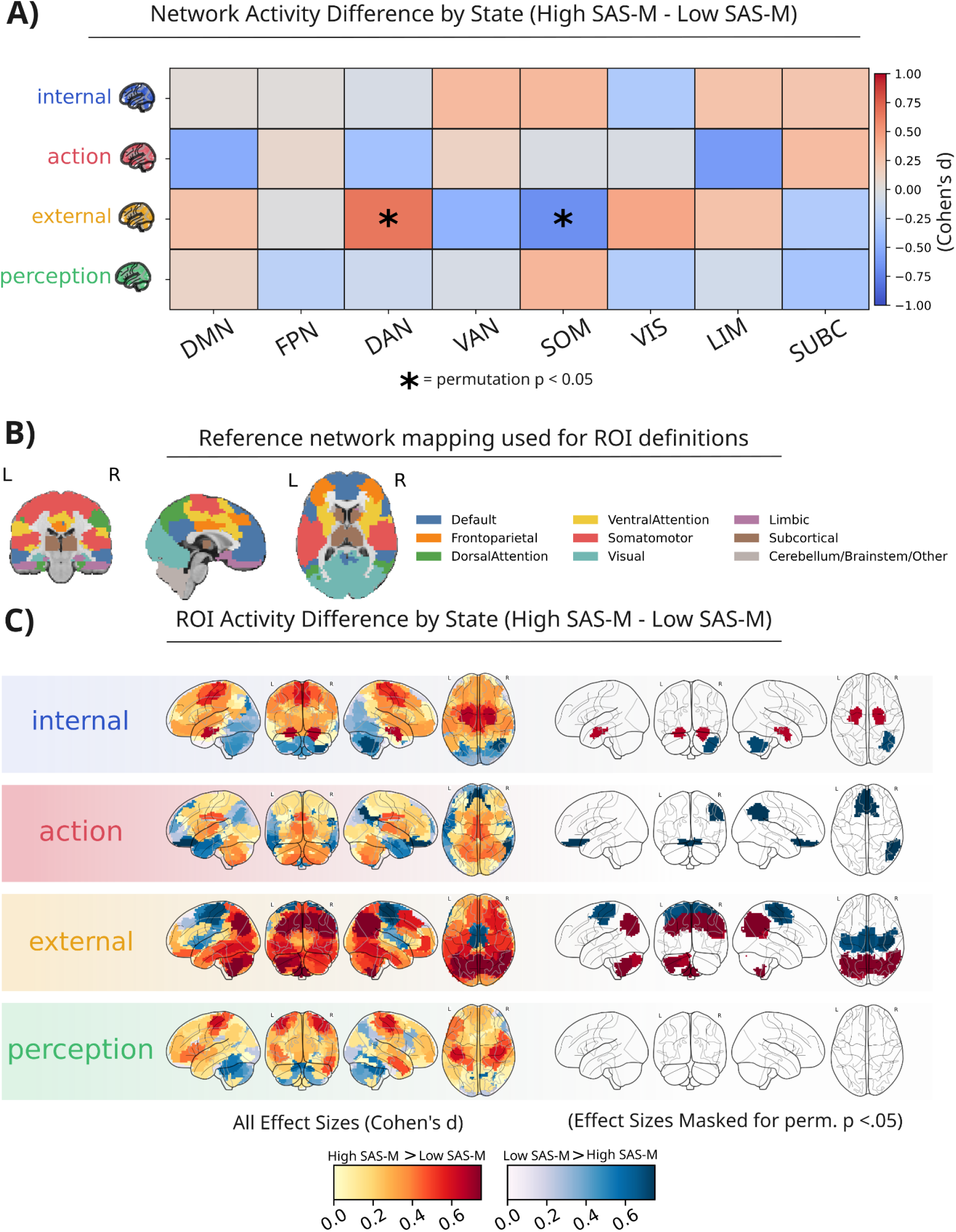
State-specific activity changes (High SAS-M and Low SAS-M) by region (networks and parcels). State-specific regional activity was averaged within subject, state, and parcel prior to group comparison. Between-group differences (High SAS-M vs Low SAS-M) were assessed using subject-level permutation tests (10,000 permutations), with Cohen’s d shown as effect size (High SAS-M minus Low SAS-M). Multiple comparisons were controlled within each state using BH-FDR. Maps are shown unthresholded and thresholded at nominal (p < 0.05) levels; no effects survived FDR correction. **A)** Network-level differences in mean BOLD activity across attractor states (internal context, action, external context, perception), shown as Cohen’s d effect sizes (High SAS-M minus Low SAS-M) for major attention networks (e.g., DMN, DAN, FPN, VIS). **B)** Approximate mapping between BASC-122 parcels and canonical large-scale networks used for network-level aggregation. **C)** Parcel-wise activity differences (High SAS-M - Low SAS-M) visualized across the cortical surface for each attractor state. Warm colors indicate greater activity in the High SAS-M group, while cool colors indicate greater activity in the Low SAS-M group. Right panels show thresholded maps (permutation p < 0.05, uncorrected). Although several regional effects reached uncorrected significance, none survived correction for multiple comparisons (BH-FDR), and results should be interpreted as exploratory.

For parcel-level analyses, group differences were tested for each of the 122 parcels using subject-level permutation tests. Specifically, the test statistic was the difference in group means, and significance was assessed using 10,000 random permutations of group labels. Effect sizes were quantified using Cohen’s d (High SAS-M minus Low SAS-M). Multiple comparisons were controlled within each state using the Benjamini-Hochberg false discovery rate (FDR) procedure.

For network-level analyses, parcels were grouped into large-scale systems using the BASC-122 network mapping (DMN, FPN, DAN, VAN, VIS, SOM, LIM, SUBC). Subject-level state-specific system activity was computed by averaging parcel values within each system. Group differences were then assessed using the same permutation testing framework and FDR correction applied across systems within each state.

For visualization, parcel-wise effect size maps were displayed both unthresholded and thresholded at nominal significance (permutation p < 0.05). FDR-corrected significance (q < 0.05) did not survive for the large number of network/parcel comparisons and thus are not displayed in Figure 4. All analyses treated subjects as independent observations and did not weight estimates by the number of TRs contributing to each state.

### 2.8 Quasi-Periodic Pattern Detection

Complex principal component analysis (cPCA) was used to identify QPP-like components from rs-fMRI data, following the framework described by Bolt and colleagues and implemented in the publicly available *complex_pca* toolbox (Bolt et al., 2022). In brief, the approach transforms preprocessed ROI time series into complex-valued analytic signals using the Hilbert transform, after which principal component decomposition is applied to the complex data matrix. This yields component time courses and complex spatial weights that jointly capture both the magnitude and relative phase structure of large-scale BOLD dynamics. Unlike conventional PCA, cPCA is specifically designed to represent time-lagged relationships among signals, allowing the recovery of propagating or anti-correlated spatiotemporal patterns rather than purely zero-lag covariance structure. Bolt et al. showed that this approach provides a compact description of resting-state fMRI dynamics and that one of the dominant components recovers QPP-like modes of the canonical task-positive/task-negative anti-correlation between DMN and DAN areas.

In the present study, cPCA was first applied at the group level for exploratory visualization and comparison by concatenating scans within each group to generate average QPPs (Supplementary Figure 3). All subsequent primary QPP analyses were conducted at the individual-subject level, with the principal network-resolved results presented in Figure 5 and supporting analyses provided in the Supplementary Material.

**Figure 5:**
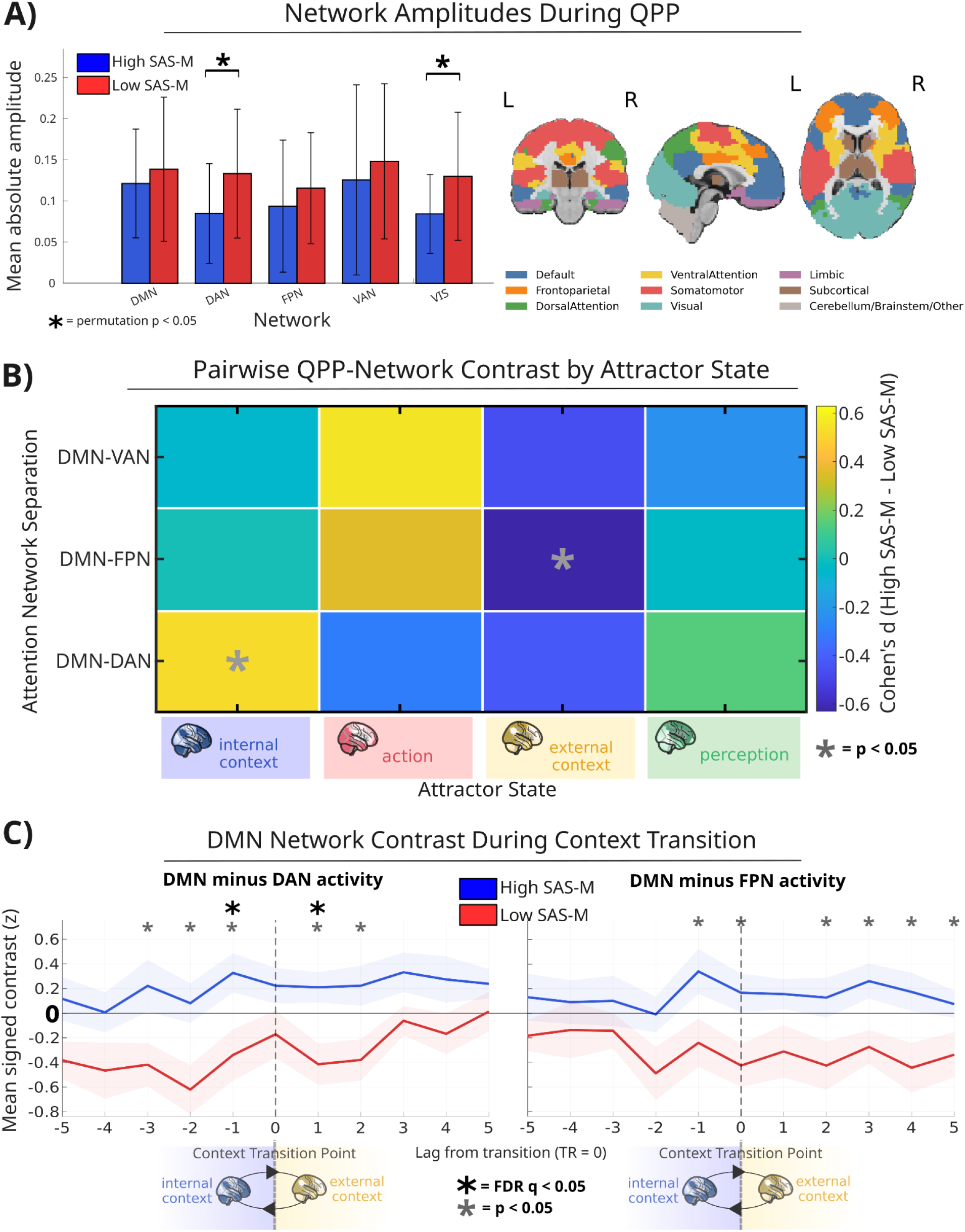
Attention Network Activity during QPPs, Attractor States, and Context Transitions. **A)** Mean absolute network amplitudes during QPP expression for major functional networks (DMN, DAN, FPN, VAN, VIS), compared between High and Low SAS-M groups. Error bars represent standard error of the mean. Asterisks denote significant group differences based on nonparametric testing. DAN and VIS were significantly lower in mean amplitude for the High SAS-M group (permutation p = 0.032 and 0.037, 10,000 permutations). **B)** Pairwise network separation (difference in QPP signal magnitude between networks) as a function of attractor state. Heatmaps display Cohen’s d effect sizes (High SAS-M - Low SAS-M) for DMN-centered contrasts (e.g., DMN-DAN, DMN-FPN, DMN-VAN) across the four attractor states (internal context, action, external context, perception). Asterisks indicate nominal significance (p<0.05, uncorrected). The High SAS-M group showed nominally increased DMN-DAN separation (p = 0.033) and decreased DMN-FPN separation (p =0.04) compared to the Low SAS-M group. **C)** Transition-triggered analysis of network dynamics during shifts along the internal-external context axis. Plots show mean signed differences in activity between DMN and attention networks (DMN-DAN, DMN-FPN) as a function of time relative to transition onset (lag in TRs). Shaded regions indicate standard error. Significant timepoints (p < 0.05 and q < 0.05 after FDR correction across lags) are marked. Results indicate increased DMN dominance relative to attention networks in the High SAS-M group during transitions between internal and external states.

### 2.9 Subject-wise Identification of QPP-like Components

Unlike prior cPCA studies performed on concatenated group-level resting-state data (Bolt et al., 2022), the present study estimated QPP-like modes of activity independently within each subject in order to preserve inter-subject variability and permit subject-level statistical inference. While the canonical DMN–task-positive QPP is consistently recovered as the dominant non-global component in concatenated datasets (i.e. component 2), the ordering and relative prominence of components varies at the individual-subject level. Accordingly, subject-wise analyses required an additional component selection procedure to identify the QPP-like spatiotemporal mode within each participant.

Consistent with prior work demonstrating that the canonical QPP is characterized by anti-correlated activity between default mode and task-positive attention systems (Bolt et al., 2022), candidate components were screened among the first three complex principal components after exclusion of globally synchronous components. The selected QPP component was defined as the non-global component exhibiting the strongest DMN-versus-DAN opposition, quantified using a combination of relative amplitude separation and phase offset across large-scale networks derived from a BASC122-to-Yeo mapping.

This procedure was intended to identify the dominant subject-specific component most consistent with the canonical DMN–task-positive QPP organization described previously using cPCA-based approaches (Bolt et al., 2022). Importantly, the selection procedure did not impose a fixed magnitude of DMN-DAN opposition or network configuration across subjects, but rather identified the non-global component exhibiting the strongest relative DMN-attention-system separation within each individual dataset. Subsequent analyses therefore quantified variability in the spatiotemporal organization, network participation, and state-dependent behavior of the selected QPP-like mode across subjects and groups.

In more detail, QPP-like mode selection focused on the first three complex principal components, which capture the majority of structured variance in resting-state data (Bolt et al., 2022). Global similarity was computed as the absolute cosine similarity between each complex component and a spatially uniform vector. Components were excluded if they were the most globally synchronous component or if global similarity exceeded 0.75. Among the remaining candidate components, the QPP score was computed as DMN-DAN opposition magnitude weighted by normalized DMN-DAN phase separation, and the highest-scoring component was selected. Components with DMN-DAN phase separation < 2.2 radians, opposition magnitude < 0.05, or composite score < 0.05 were classified as weak; components with no eligible candidate were classified as absent. In the present dataset, all subjects exhibited identifiable QPP or weak-QPP components (Supplementary Figure 1).

### 2.10 QPP Sign Alignment for Network Analysis

Because complex principal components are defined up to an arbitrary sign, the orientation of QPP components can vary across subjects, leading to apparent inversions of network activity (e.g., DMN up vs. down) that are not meaningful. This is another consideration not encountered when running cPCA on concatenated group-level scans (such as in Bolt et al., 2022). To ensure network interpretability across subjects, we sign-aligned QPP components prior to analysis. For subject-level analyses, the selected QPP component was oriented such that the DMN exhibited consistent polarity across participants. This alignment ensured that comparisons of network phase relationships and anticorrelation structure (e.g., DMN vs. attention networks) reflected true differences in spatiotemporal dynamics rather than arbitrary sign differences.

### 2.11 Exploratory Subject-Wise QPP-Attractor Analyses

To explore the temporal relationship between whole-component QPP expression and attractor-state organization, a subject-specific QPP time course was derived independently for each participant from the selected cPCA component. The real component score was extracted at each TR, sign-aligned to maintain consistent DMN polarity across participants, z-scored, and temporally matched to the corresponding attractor-state labels. Example time courses and group-averaged network trajectories are presented descriptively in Supplementary Figure 4 to illustrate the relative timescales of continuous QPP fluctuations and discrete attractor-state transitions.

Exploratory subject-level analyses additionally tested whether attractor states were differentially represented at QPP peaks, troughs, and zero-crossings and characterized signed QPP expression surrounding transitions along the internal–external context and action–perception axes. Between-group differences in these measures were evaluated using two-sided nonparametric tests and are reported in Supplementary Figure 2.

### 2.12 Subject-wise QPP network behavior vs attractor states

To characterize network-resolved QPP dynamics across attractor states (Figure 5), we analyzed QPPs estimated independently for each subject from their selected subject-specific cPCA component. For each subject, the reconstructed ROI-by-TR matrix of the selected component was loaded, sign-aligned so that DMN polarity was consistent across subjects, and then averaged within Yeo7-derived networks of interest: DMN, DAN, FPN, VAN, and VIS. Whole-scan network amplitude was quantified as the mean absolute value of each subject’s network-specific reconstructed QPP time course across the full scan. Importantly, these measures reflect the magnitude of network participation within the QPP component itself, rather than raw BOLD signal amplitude. Between-group differences in these amplitude measures were assessed using both Wilcoxon rank-sum tests and subject-level permutation tests on the difference in means (10,000 permutations), with Cohen’s d reported as the effect size.

To examine context-dependent inter-network relationships, DMN-centered pairwise contrasts were computed at each TR as simple within-timepoint differences between the mean reconstructed network signals (DMN-DAN, DMN-FPN, DMN-VAN). For the state-wise analyses, absolute values of these contrasts were taken to quantify network separation irrespective of sign, then z-scored within-subject and averaged within each attractor state. Group comparisons of state-wise absolute separation were performed using Wilcoxon rank-sum tests, and permutation p-values were additionally computed for descriptive robustness. Cohen’s d (High SAS-M - Low SAS-M) was used for the heatmap effect sizes. Because these comparisons were exploratory and none of the nominal state-wise effects survived correction, the panel was displayed with raw rank-sum significance markers while permutation p-values were retained for reporting.

To assess network dynamics around attractor transitions, signed DMN-centered contrasts were z-scored within-subject and averaged in a ±5 TR window around transition events. Transition-triggered analyses were restricted to the two theoretically relevant transition classes: internal-external context transitions and action-perception transitions. Between-group comparisons were performed at each lag using Wilcoxon rank-sum tests, and Benjamini-Hochberg FDR correction was applied across lags within each network pair and transition type. In the transition plots, raw nominal p < 0.05 timepoints were marked separately from FDR-significant lags.

## 3. Results

### 3.1 Qualitative Differences in Attractor State Space

To visualize large-scale attractor organization, subject trajectories were projected into a two-dimensional space defined by internal–external context and perception–action dimensions (Figure 2A). Both groups exhibited similar state-space coverage, with the greatest density of time points near the origin. Subject-level mean positions are shown at an expanded scale in Figure 2B. Both group centroids remained close to the origin, although the High SAS-M group showed a modest descriptive shift toward the internal-context (left quadrant, Figure 2A) and action directions (upper quadrant) and greater dispersion (Figure 2B). These visual differences were descriptive and were not interpreted as evidence of statistically reliable group separation.

### 3.2 Attractor State Dynamics

Subject-level comparisons of attractor-state occupancy did not reveal robust group differences. An omnibus permutation test comparing four-state occupancy vectors between groups was not significant (p = 0.454; 10,000 permutations), and no subject-level Mann-Whitney U comparison reached nominal significance for state fractions, axis scores, mean dwell times, state entropy, or switch rate (all p > 0.05).

The largest non-significant trend was for external-context fraction, which was lower in the High SAS-M group than the Low SA group (means 0.225 vs 0.276, p = 0.121, Cohen’s d = −0.48, Cliff’s delta = −0.29). Mean external-context dwell time showed a similar trend (p = 0.071, d = −0.54), whereas mean action dwell time showed only a modest effect (p = 0.308, d = 0.39). State entropy and switch rate were closely comparable between groups (p = 0.455 and 0.744, respectively), arguing against a strong group difference in overall switching dynamics or attractor-state diversity. Levene’s test suggested unequal variance only for action dwell time (p = 0.036), while all other variance comparisons were non-significant.

### 3.3 Attractor States vs Questionnaire Scores

Exploratory associations of SAS-M and DASS-21 scores with subject-level attractor and QPP-derived measures yielded several nominally significant relationships; however, none survived correction for multiple comparisons (Supplementary Table 1). These associations were therefore not interpreted further. See discussion for more on self-report / questionnaire measures.

### 3.4 Regional activity by attractor state

State-specific regional activity differences between High and Low SA groups were modest and spatially heterogeneous. At the network level, permutation testing identified a small number of nominally significant effects (permutation p < 0.05), primarily within the external-context state, including increased dorsal attention network activity and reduced somatomotor activity in the High SAS-M group. However, none of these effects survived FDR correction across networks within-state. Parcel-wise analyses revealed distributed patterns of moderate effect sizes across multiple states, particularly in the internal-context, action, and external-context states. Several regions reached nominal significance (p < 0.05), but no parcel survived FDR correction within any state, and the results should be considered exploratory.

These results suggest that while state-dependent regional activity differences exist between groups, they are subtle, spatially distributed, and not robust to multiple-comparison correction, consistent with a diffuse rather than focal BOLD signature of attention network changes.

### 3.5 Exploratory Group-Level QPP Network Analysis

Exploratory group-template reconstructions qualitatively suggested greater DMN-DAN and DMN-FPN phase separation in the High SAS-M group relative to the Low SAS-M group (Supplementary Figure 3). Because these phase relationships were derived from group-level cPCA templates and were not reproduced as significant differences in the subject-level analyses, they are presented descriptively and are not interpreted as inferential group effects.

### 3.6 Whole-Component QPP Expression Across Attractor States

Using subject-specific cPCA-derived QPPs, we examined whether whole-component QPP expression differed across attractor states or between SAS-M groups. Although QPP expression varied across time and attractor states within individual participants, subject-level summaries did not reveal consistent differences between groups (Supplementary Figure 2). These findings motivated subsequent analyses examining whether group differences were more specifically localized to network-resolved and transition-related aspects of QPP dynamics.

### 3.7 QPP Network-Level Expression vs Attractor States and Transitions

To further characterize how large-scale networks contribute to QPP dynamics across cognitive contexts, we examined network-level QPP expression and inter-network relationships as a function of attractor state (Figure 5). Network-resolved analyses of subject-specific QPPs revealed clearer group differences than observed at the whole-component level. High SAS-M subjects showed reduced QPP amplitude in externally oriented networks, most notably in the dorsal attention network (DAN; rank-sum p = 0.0187; permutation p = 0.0325; *d* = −0.68), with a similar effect in the visual network (permutation p = 0.0370; *d* = −0.69). No differences were observed for DMN, FPN, or VAN. These amplitude measures reflect network participation within the reconstructed QPP component, rather than overall BOLD activity, indicating reduced engagement of DAN and visual systems specifically within QPP dynamics.

State-wise analyses of absolute DMN-centered network separation showed modest, context-dependent effects. DMN-DAN separation was greater in High SAS-M during the internal-context state (p = 0.0328, *d* = 0.52), while DMN-FPN separation was reduced during the external-context state (p = 0.0401, *d* = −0.63). However, these effects were not supported by permutation testing and did not survive multiple-comparison correction, and are therefore interpreted as exploratory.

The most consistent group differences emerged in transition-triggered analyses of signed network contrasts during internal↔external context transitions. For DMN-DAN, High SAS-M subjects showed greater DMN dominance around transitions, with FDR-corrected significant effects at lag −1 (p = 0.0083, q = 0.0457) and lag +1 (p = 0.0018, q = 0.0197), indicating sustained DMN elevation relative to DAN both before and after transitions. DMN-FPN showed weaker, non-corrected effects, and no differences were observed for DMN-VAN or during action↔perception transitions.

Importantly, reduced DAN involvement here is specific to QPP dynamics and does not reflect reduced overall DAN activity. Prior analyses, using attractor states only (Figure 4), showed nominally increased DAN activity during externally oriented states, suggesting a dissociation between baseline state engagement and participation in large-scale quasi-periodic dynamics, with High SAS-M characterized by reduced integration of attention networks into the dominant QPP mode. Among the analyses performed, the most robust effects were observed specifically during transitions between internal and external attractor states, where High SAS-M subjects consistently showed greater DMN dominance relative to attention networks (Figure 5C).

## 4. Discussion

The present findings suggest that differences associated with higher smartphone addiction scores were more apparent in large-scale spatiotemporal network dynamics than in attractor-state occupancy. Across analyses, the clearest effects emerged in the network-resolved QPP results, particularly in the altered participation of attention-related systems and increased DMN dominance around internal-external context transitions. In contrast, attractor-state occupancy and subject-level summary metrics were broadly similar between groups.

### 4.1 Attractor State Summary

Analysis of attractor-state dynamics revealed no robust group differences in overall state occupancy or subject-level summary metrics. Nevertheless, qualitative examination of the attractor-state space suggested a tendency for individuals with higher SAS-M scores to occupy regions associated with action-oriented states and to exhibit broader dispersion within the attractor landscape. The state-specific activities by region (Figure 4) revealed strong trends for elevated engagement in regions such as occipital visual areas during external states and bilateral amygdala in internal states (Figure 4C), but these trends did not survive multiple comparisons correction.

### 4.2 Group-Level QPP Summary

At the group level, separately reconstructed QPP templates qualitatively showed greater apparent phase separation between the DMN and both DAN and FPN in the High SAS-M group (Supplementary Figure 3). These group-template patterns are broadly consistent with prior work linking increased segregation between default-mode and task-positive systems to task engagement (Watters et al., 2025; Seeburger et al., 2026). However, because group-template reconstructions do not provide independent subject-level estimates and the corresponding differences were not consistently reproduced in subject-level analyses, they are presented descriptively and should be interpreted cautiously.

### 4.3 Subject-Specific-QPP Component vs Attractor States

Whole-component QPP expression did not show reliable group differences across attractor states. This provides limited support for a generalized alteration of QPP-attractor coupling and instead suggests that group effects, where present, are more specific to network-resolved or transition-dependent aspects of QPP dynamics.

### 4.4 QPP Network Expression by Attractor State

The clearest and most statistically robust findings emerged during transitions between internal and external attractor states, where individuals with higher smartphone addiction scores showed increased DMN dominance relative to the dorsal attention network (DAN) during subject-specific QPP dynamics. These effects survived correction for multiple comparisons and were evident both immediately before and after context transitions, suggesting altered coordination between internally and externally oriented attention systems during periods of dynamic state reorganization. Similar but weaker trends were observed for DMN-frontoparietal (FPN) relationships, whereas no consistent differences were observed for ventral attention (VAN) dynamics or action-perception transitions.

Beyond transition-related effects, network-resolved analyses also showed reduced contribution of externally oriented networks within the reconstructed QPP mode itself, particularly in the DAN and, to a lesser extent, the visual network (VIS). In addition, exploratory state-wise analyses suggested increased DMN-DAN separation during internally oriented states and reduced DMN-FPN separation during externally oriented states, although these effects did not survive permutation testing or correction for multiple comparisons. Taken together, the results suggest that the primary alterations associated with higher smartphone addiction scores may involve disrupted coordination of attention networks during transitions between cognitive contexts, rather than large static shifts in attractor-state occupancy or global QPP expression.

### 4.5 Synthesis of Results

Together, the present findings suggest that smartphone addiction is associated more strongly with altered attention-network dynamics during cognitive-state transitions than with large differences in general resting-state dynamics. The clearest effects emerged during transitions between internal and external attractor states, where individuals with higher smartphone addiction scores showed increased DMN dominance relative to attention-related systems, particularly the DAN, during subject-specific QPP dynamics. Complementary analyses additionally suggested reduced participation of externally oriented networks within the reconstructed QPP mode itself, along with exploratory alterations in state-dependent DMN-DAN and DMN-FPN relationships.

Several of these features, particularly increased DMN-attention-network separation, resemble patterns previously reported during task-engaged or externally directed cognitive states in QPP studies (Seeburger et al., 2024; Watters et al., 2025). Recent work combining QPP analysis with task fMRI further demonstrated that decoupling between the DMN and FPN is associated with elevated cognitive demand during working-memory tasks (Seeburger et al., 2026). In the present study, qualitatively similar patterns were observed during resting-state activity in High SAS-M subjects, including increased DMN-FPN separation and altered DMN-attention-network relationships. Although these effects were modest and not uniformly robust across analyses, their presence during rest may suggest subtle differences in baseline coordination between internally and externally oriented systems during spontaneous activity.

The attractor transition-related findings may further indicate altered coordination between internally and externally oriented attentional systems during periods of cognitive-state reorganization. Within the broader framework of attentional control theory (Eysenck & Derakshan, 2011), efficient attentional shifting depends on the limited resources of executive control networks, i.e. networks such as the FPN and DAN that support top-down attentional regulation and goal-directed processing via interactions with the DMN (Dixon et al., 2018). Although attentional control theory was originally developed in the context of anxiety-related attentional bias (Eysenck & Derakshan, 2011), the broader framework may also be relevant to persistent high-stimulation digital engagement, where frequent switching between salient stimuli could place sustained demands on executive control systems. In this context, increased DMN dominance relative to the DAN and FPN during internal-external attractor transitions may reflect reduced efficiency in reallocating attentional resources during moments of shifting cognitive context. However, these interpretations remain speculative and cannot be directly inferred from resting-state data alone. Future work combining task-based paradigms, objective smartphone-use measures, and dynamic network analyses will be necessary to determine whether altered QPP-attractor dynamics are associated with measurable impairments in attentional control or executive flexibility.

Given the modest sample size and exploratory nature of several analyses, these findings should be interpreted as hypothesis-generating. Nevertheless, they provide a preliminary framework for understanding how habitual engagement with high-stimulation digital environments may relate to alterations in intrinsic large-scale brain dynamics. Future work with larger samples and longer acquisitions will be necessary to determine whether these effects represent stable trait-level differences or context-dependent adaptations to patterns of digital engagement.

### 4.6 Limitations

Several limitations should be considered when interpreting the present findings. First, head motion was addressed through exclusion of high-motion participants in the original dataset (Abdul Rashid et al., 2021) and regression of motion parameters during preprocessing in the CONN pipeline. Remaining subjects did not differ significantly in framewise displacement between groups. Nevertheless, residual motion effects cannot be fully excluded and may subtly influence dynamic fMRI measures.

Second, the sample size was modest (N = 40), limiting statistical power and likely contributing to the observation that several moderate to strong effects did not survive correction for multiple comparisons. Neuroimaging measures, especially functional correlation, are notoriously susceptible to poor replicability and generalizability in small sample sizes (Marek & Laumann, 2025). While we applied dynamic measures such as fcANN that show robustness across datasets (Englert et al., 2025) these findings are susceptible to the same fMRI pitfalls and should be interpreted as exploratory and hypothesis-generating rather than definitive evidence of group differences.

Third, QPP detection at the individual-subject level was performed on relatively short resting-state scans, which may reduce the stability and reliability of estimated spatiotemporal patterns. This highlights the need for longer acquisitions or replication in larger datasets. More broadly, while subject-specific identification of QPP-like components may provide a useful framework for studying inter-individual variability in large-scale spatiotemporal dynamics, the reliability and optimal selection criteria for individual-level cPCA-derived QPP estimation remain areas for further methodological validation.

Fourth, while attractor-based and QPP approaches provide complementary insight into dynamic brain organization, both rely on modeling assumptions (e.g., predefined attractor embeddings, component selection criteria in cPCA) that may influence results. Future work using alternative dynamic analysis frameworks would help establish the robustness and generalizability of the observed effects. In addition, the cross-sectional design precludes causal inference regarding the directionality of the relationship between smartphone use and brain dynamics.

Fifth, smartphone addiction was assessed using self-report questionnaire measures, which may be subject to reporting bias and may not fully capture actual usage behavior (Coyne et al., 2023). Objective measures of smartphone use (e.g., screen time logs, app usage metrics) would provide a more precise characterization of digital engagement and its relationship to brain dynamics.

Finally, as with the original study of this dataset, no significant group differences in age or sex distribution were identified, although the sample size limits excluding demographics as a confound. Future studies with larger samples and objective behavioral measures will be important for evaluating the specificity of the reported effects (see below).

### 4.7 Smartphone Use as a Consideration in Neuroimaging Studies

Objective measures of smartphone use are rarely collected or reported in neuroimaging datasets of smartphone addiction, often relying on self-report/questionnaire data instead (Korte, 2020). The present findings, together with prior work, suggest that excessive smartphone use may contribute to unmeasured heterogeneity in fMRI studies more broadly, beyond studies specifically focused on behavioral addiction. Given the growing body of behavioral and neuroimaging evidence linking problematic smartphone use to alterations in attention, cognitive control, and cortical network organization (Liebherr et al., 2020; Anbumalar & Binu Sahayam, 2024), it may be important to consider smartphone usage as a standard demographic variable in future neuroimaging studies.

Large scale consortia neuroimaging efforts such as the Human Connectome Project and UK Biobank already document demographic and behavioral data on age, sex, intelligence, sleep, substance use, psychiatric history, etc. (Glasser et al., 2016; Miller et al., 2016). Incorporating objective measures such as screen time, social media time, or interaction frequency into large-scale consortia efforts could help clarify the brain-behavior effects of smartphone addiction. This is especially important given the aforementioned precarity of small sample neuroimaging studies, which may require sample sizes into the hundreds before small effects and correlations are replicable and generalizable (Marek & Laumann, 2025). Given the need for longer scan times and higher sample sizes (Ooi et al., 2025), small-scale studies focused exclusively on smartphone addiction may face substantial challenges in achieving the sample sizes needed for robust and generalizable brain-behavior inference.

## 5. Conclusions and Future Directions

In summary, the clearest effects observed in the present study involved altered coordination between default mode and attention networks during transitions between internally and externally oriented attractor states. Individuals with higher smartphone addiction scores showed increased DMN dominance and reduced participation of attention-related networks, particularly during subject-specific QPP dynamics. Although several findings were exploratory or modest in isolation, a pattern across the complementary dynamic fMRI analyses suggested altered state-dependent organization of large-scale resting-state brain dynamics in smartphone addiction. This is consistent with evidence that brain network dynamics may serve as transdiagnostic and dimensional biomarkers of psychiatric and psychological disorder (Cocuzza et al., 2026).

These findings are preliminary and hypothesis-generating, but support further investigation of the relationship between excessive smartphone use and resting-state brain dynamics and the coordination of attention networks. Future work incorporating longitudinal designs, objective smartphone usage metrics, and task-based paradigms will be critical for establishing causality and clarifying the functional significance of these effects.

As digital technologies continue to shape daily cognitive experience, understanding their relationship to intrinsic brain dynamics will be increasingly important for both basic neuroscience and mental health research.

## Code and Data Availability

Attractor network dynamic analysis used the publicly available code from Englert and colleagues (Englert et al., 2025, repository: https://github.com/pni-lab/connattractor). Complex principal component analysis was performed using the publicly available implementation released by Bolt and colleagues (repository: https://github.com/tsb46/complex_pca) for fMRI analyses (Bolt, 2022). The open-source SPM-based functional connectivity toolbox (CONN) was used in MATLAB for all preprocessing (Whitfield-Gabrieli, S., and Nieto-Castanon, 2012). Nilearn tools were used for interaction with the Connattractor toolbox and some data visualization (https://nilearn.github.io). All additional code used for preprocessing, analyses, and figure generation was developed in-house using Python 3.14 (Python Software Foundation) and MATLAB R2024a (MathWorks, Natick, MA, United States). The main analysis code is available at: https://github.com/Dalimear/smartphone-qpp-cPCA-attractor-dynamics. The raw or preprocessed fMRI scans may be made available upon reasonable request from the original study authors.

## Acknowledgments

This study was made possible by collaboration with the original scan acquisition and study authors: Aida Abdul Rashid, PhD, and Subapriya Suppiah, MD, and others (Abdul Rashid et al., 2021) at the Universiti Putra Malaysia (UPM). We also thank Mr. Mohd Khalil Saleh, chief technologist at the Section for Nuclear Medicine, Department of Radiology, Hospital Sultan Abdul Aziz Shah, UPM for his diligence in making the data available in the local institutional repository. The study was also supported by the project “Research of Excellence on Digital Technologies and Wellbeing CZ.02.01.01/00/22_008/0004583” which is co-financed by the European Union.

The authors also acknowledge the use of Claude (Anthropic, Claude Sonnet 4.6) and ChatGPT (OpenAI, GPT-5.5) for code assistance/debugging and late-stage manuscript grammar editing and typo identification. Final code was reviewed for correctness. All literature review, citation, bibliography, drafting, scientific interpretation, study design, analysis, etc., was done by the authors. No AI-generated figures or images were used. All figures were produced and assembled manually using open source vector graphics software (Inkscape Project, https://inkscape.org).

## Supplementary

**Supplementary Table 1:**
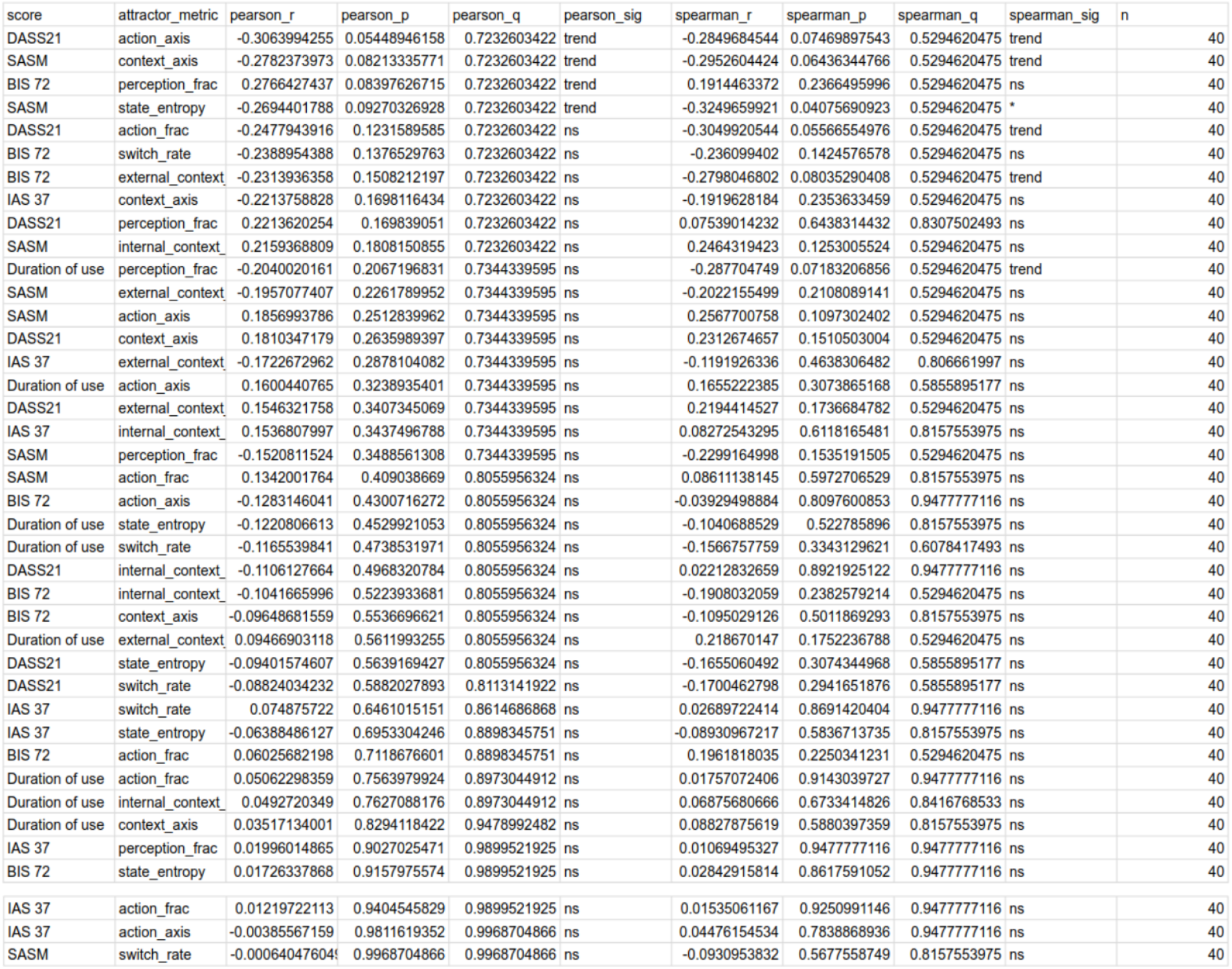
Summary of Attractor State Metrics vs Subject Behavioral Scores. Note that while some trends were observed with respect to smartphone addiction and depression-anxiety scores, no correlations reached corrected significance. No other metrics vs behavioral score correlations survived correction for multiple comparisons.

**Supplementary Figure 1:**
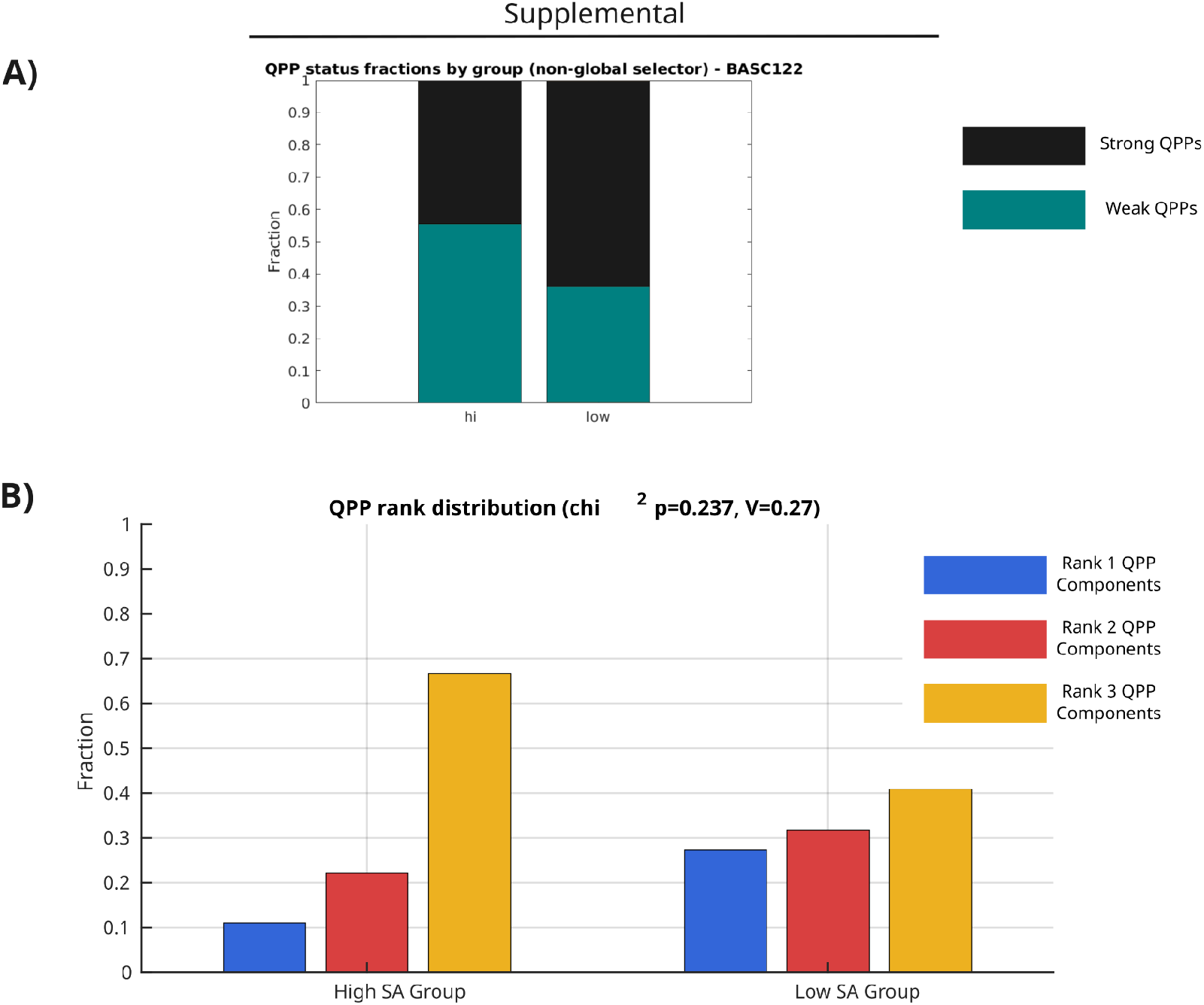
**A)** Fraction of strong vs weak QPPs. Note, all subjects had at least weak QPPs. **B)** Distribution of QPP components by rank (from complex PCA). Note the trend towards lower rank (and thus less explained variance) in the High SAS-M group. A tendency towards ‘weaker’ QPPs (less network opposition and amplitude) may contribute to the observed phenomenon of reduced DAN and VIS network amplitude during QPPs in the High SAS-M group.

**Supplementary Figure 2:**
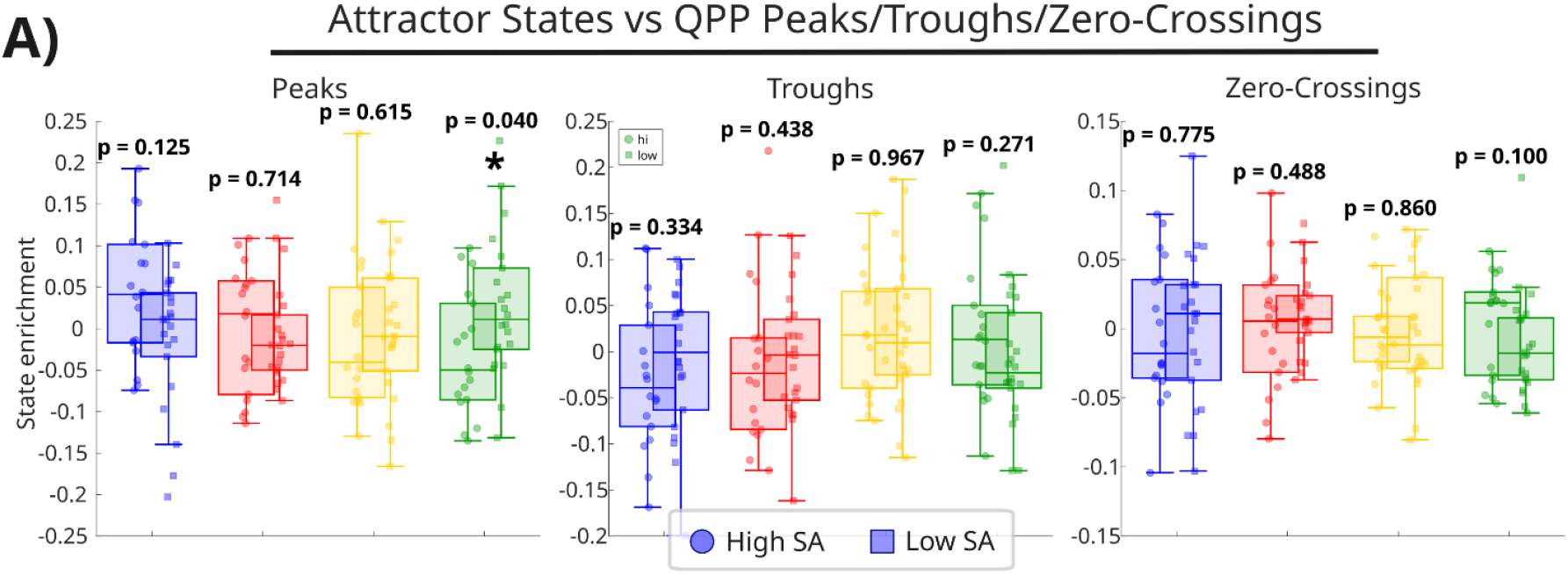
Extra Whole-QPP vs attractor state metric summary. **A)** QPP phases (peaks/troughs/zero-crossings) by attractor states.

**Supplementary Figure 3:**
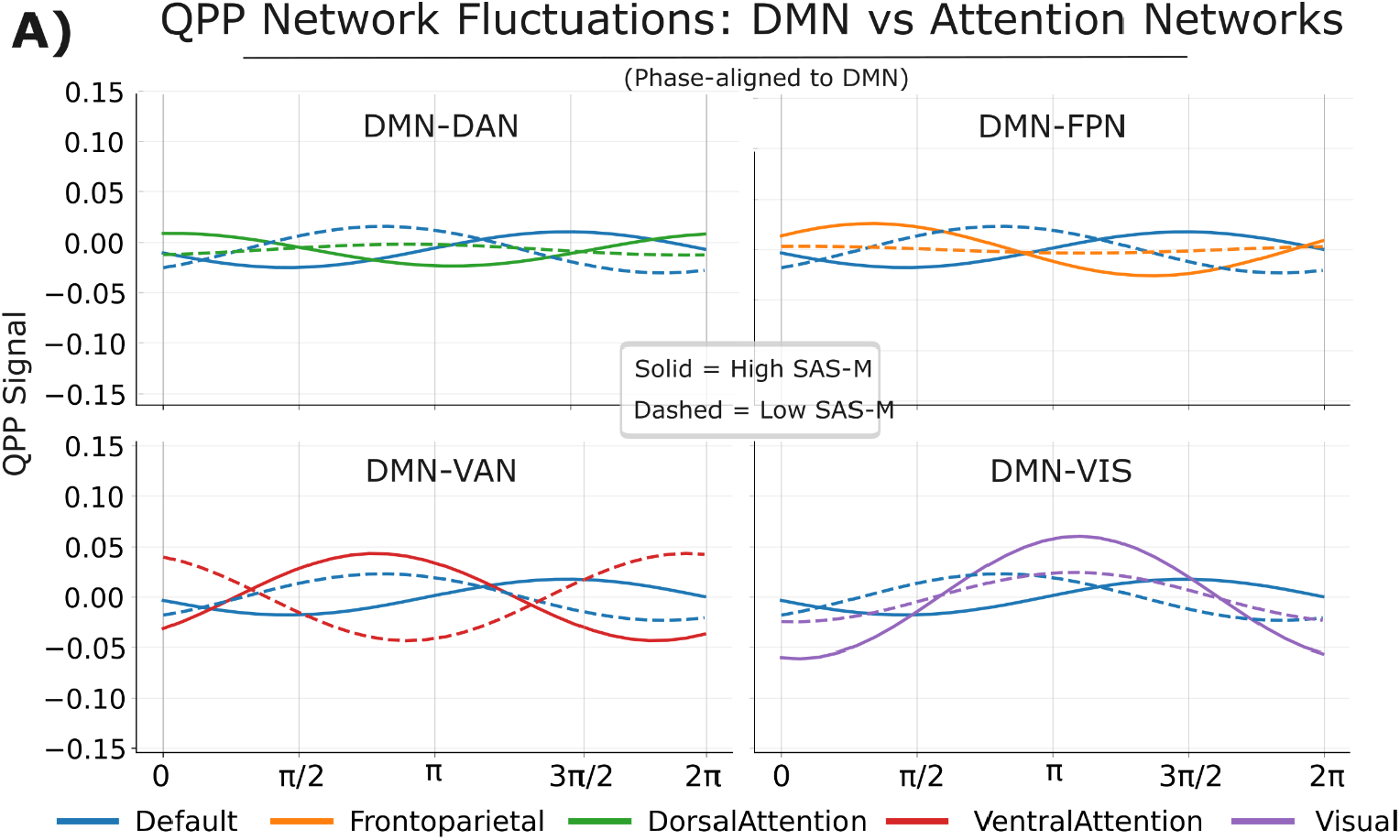
Exploratory group-level QPP network phase relationships. Network trajectories were reconstructed from separate group-level cPCA templates and phase-aligned to the DMN within each group. The High SAS-M template qualitatively showed greater DMN-DAN and DMN-FPN phase separation than the Low SAS-M template. Because these trajectories are derived from group-level templates rather than independent subject-level estimates, the figure is presented descriptively and no inferential statistics are reported.

**Supplementary Figure 4:**
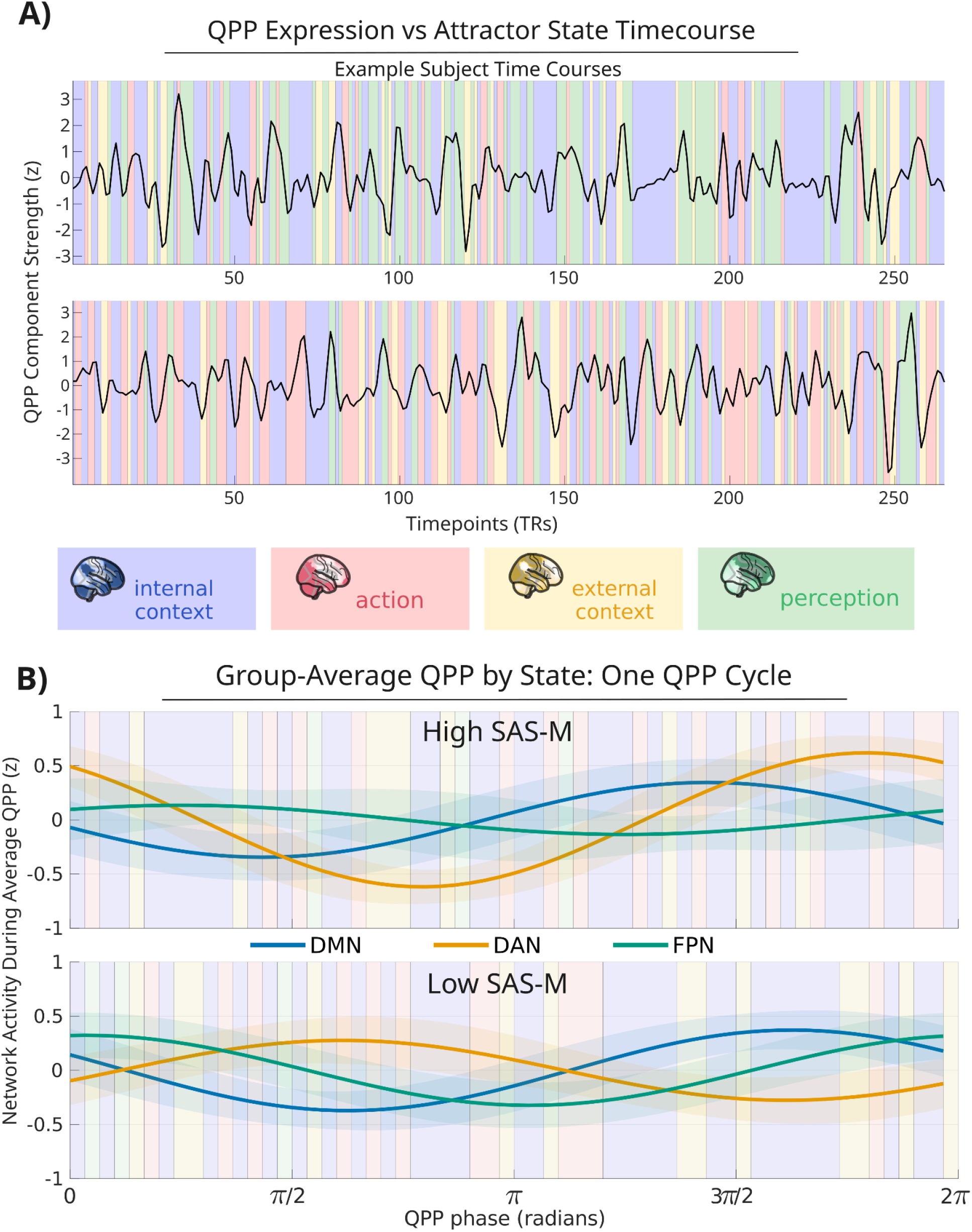
Descriptive visualization of QPP expression relative to attractor-state dynamics. Subject-specific QPP expression was derived from each participant’s selected cPCA component, sign-aligned across participants, and temporally matched to attractor-state labels at each TR. The figure is intended to illustrate the relative timescales of discrete attractor-state transitions and continuous infraslow QPP fluctuations; subject-level analyses did not identify significant between-group differences in whole-component QPP expression across attractor states. **A)** Example participant time courses showing z-scored QPP component expression overlaid with attractor-state labels. Colored background segments indicate internal context (blue), action (red), external context (yellow), and perception (green), illustrating that attractor-state assignments can change multiple times over the course of an infraslow QPP fluctuation. **B)** Group-averaged QPP network trajectories across one phase cycle (0–2π), shown separately for High and Low SAS-M groups. Lines represent mean signed contributions from the default mode (DMN), dorsal attention (DAN), and frontoparietal (FPN) networks, with shading indicating ±SEM across participants. Background colors indicate the dominant attractor-state assignment at each QPP phase. Panel B is provided as a descriptive visualization of the temporal relationship between network-level QPP expression and attractor-state organization; no significant between-group differences in whole-component QPP-state coupling were identified.

